# Mutational effects of ubiquitously present gamma radiation on *Arabidopsis thaliana*: insight into radiosensitivity in the reproductive stage

**DOI:** 10.1101/2021.02.03.429546

**Authors:** Akira S. Hirao, Yoshito Watanabe, Yoichi Hasegawa, Toshihito Takagi, Saneyoshi Ueno, Shingo Kaneko

## Abstract

Earth has always been exposed to ionizing radiation from natural sources, and man-made sources have added to this radiation. In order to assess mutational effects of ubiquitously present radiation on plants, we performed a whole-genome resequencing analysis of mutations induced by chronic irradiation throughout the life-cycle of *Arabidopsis thaliana* under controlled conditions. Resequencing data from 12 M_1_ lines and 36 M_2_ progeny derived under gamma-irradiation conditions ranging from 0.0 to 2.0 Gy/d were obtained to identify *de novo* mutations, including single base substitutions (SBSs) and small insertions/deletions (INDELs). The relationship between *de novo* mutation frequency and a low-to-middling dose of radiation was assessed by statistical modeling. The increasing of *de novo* mutations in response to doses of irradiation fit the negative binomial model, accounting for the high variability of mutation frequency observed. Among the different types of mutations, SBSs were more prevalent than INDELs, with deletions being more frequent than insertions. Furthermore, we observed that the mutational effects of chronic radiation are more intensive during the reproductive stage. These outcomes could provide valuable insights into practical strategies for environmental radioprotection of plants on Earth and in space.

## INTRODUCTION

Since the discovery of Muller (1927) [1] and Stadler (1928) [2] that ionizing radiation (X-rays) induces mutations, the biological effects of ionizing radiation have been extensively studied in the field of genetics. In the last nearly one hundred years, the mutagenesis properties of ionizing radiation have been most intensively studies over short durations and at acutely high doses, while few controlled experiments have used long-term or chronic low doses of irradiation, although field research under chronic low doses of radiation were often conducted for environmental radioprotection [reviewed in 3, 4]. Furthermore, current relevant knowledge of radiological mutagenesis is mostly derived from humans and other animals rather than plants, although animals and plants have significant differences in radiobiology. For example, a significantly higher dose of ionizing radiation is typically required to impair plant cells as compared to animal cells [5]. These contextual issues as well as the impacts of chronic irradiation remain to be studies in higher plants, especially in controlled conditions.

As whole-genome sequencing technologies advance rapidly, genome-scale surveys for irradiated plants have been performed to characterize the properties and frequency of mutations [6–14, reviewed in 15]. Most of these genome-scale studies have stemmed from mutation breeding, and thus acute irradiation experiments were performed to identify novel mutations and characterize mutation profiles and the effectiveness/efficiency of radiological mutagens, such as gamma-ray and carbon ion beams. As an exceptional study, Hase et al. (2020) conducted chronic gamma irradiation studies with 0.002–500 mGy/h in five successive generations of *Arabidopsis thaliana*, and found that mutation frequency did not increase, but rather the ratio of transition to transversion mutations decreased at very low dose rates (~1 mGy/h), suggesting complementary DNA repair activities [14]. In the context of environmental radioprotection, however, Hase et al. (2020) did not completely reveal the effects of ubiquitously present radiation, as their chronic irradiation was performed on plants in the vegetative growth stage (approximately two weeks before bolting), but not in the reproductive stage. Generally, the reproductive growth stage is more radiosensitive than the vegetative stage, although life stage-specific sensitivity to mutations has not been explored on a genome-wide scale. In this study, therefore, we performed a whole-genome resequencing analysis of mutations induced by life cycle-specific chronic irradiation in *A. thaliana*.

Earth has always been exposed to ionizing radiation from natural sources, and man-made sources have added to this radiation. Absorbed dose rates in the air of terrestrial gamma radiation typically range from 10 to 200 nGy/h over 55 countries worldwide [16]. Another field where the effects of ubiquitously present radiation are important is space, where irradiation dose rates are more than a hundred times higher than in terrestrial environments. For example, the radiation dose at the International Space Station (ISS) can vary, but is estimated to be on average 0.327 mGy/d [17]. From the perspective of radioprotection, a ‘low dose’ of radiation from any man-made sources has been defined as 100 mGy or less of sparsely ionizing radiation, and a ‘low dose-rate’ as less than 0.1 mGy/min of sparsely ionizing radiation when averaged over about one hour [18]. Although few studies have examined the effects of chronic low dose ionizing radiation on plants, the United Nations Scientific Committee on the Effects of Atomic Radiation (UNSCEAR) (1996) suggests 10 mGy/d as a threshold dose rate for environmental radioprotection of plants, even with some sensitive difference among species [19]. For more detailed examples, the International Commission on Radiological Protection (ICRP) (2004) provides Derived Consideration Reference Levels (DCRLs) for grass (1-10 mGy/d) and for pine trees (0.1-1 mGy/d) [20]. Additionally, as mentioned above, the mutational effect of ionizing radiation at very low dose rates (~1 mGy/h) has not been determined in plant models [14]. Thus, the mutational effect of ‘low doses’ and/or ‘low dose-rates’ above the natural background radiation are difficult to reliably distinguish between in terms of low risk or zero risk. In this study, therefore, we used low-to-moderate doses of gamma radiation (less than 2 Gy/d) to more clearly identify the mutational effects of chronic radiation throughout the life-cycle of plants.

Mutations are intrinsically difficult to study because *de novo* mutation events are very rare, even under exposure to ionizing radiation. In order to overcome the rarity of mutation events, mutation accumulation lines have been used as a principal way to estimate the rates and properties of mutations [21, 22, reviewed in 23]. The increase in mutations accumulating with the advancement of generations is useful to accurately estimate mutation profiles, but which are not the same at the occurrence of the mutation, because germline *de novo* mutations are allowed to drift to fixation in inbred lines. In mutation accumulation lines of self-fertilizing organisms such as *A. thaliana* and *Oryza sativa*, only approximately half of *de novo* mutations will be fixed as homozygous mutations in later generations due to genetic drift. Thus, as an alternative approach, identifying *de novo* germline mutations in M_2_ generation, i.e., the subsequent generation immediately after mutagenesis treatment, is worthwhile for assessing the *de novo* mutation profile before successive genetic drift. *De novo* mutations that occur before gamete formation are transmitted to selfed progeny as homozygous or heterozygous variants according to the Mendelian segregation ratio of 1:2. On the other hand, mutations that occur after gametogenesis must be observed as heterozygous in the M_2_ generation. Thus, if the mutational effects of chronic radiation were more intensive in the reproductive growth stage than in the vegetative growth stage, an excessive number of heterozygous mutations above the Mendelian segregation ratio would be observed in the M_2_ generation. Therefore, in this study, we applied the latter approach to identify *de novo* mutations in M_2_ generation to test the hypothesis that radiation-induced mutations more frequently occur in the reproductive stage than in the vegetative stage.

The main objective of the present study was to assess the mutation frequency and spectrum induced by chronic irradiation throughout the life-cycle of the model plant, *A. thaliana*. Resequencing data from 12 M_1_ lines and 36 M_2_ progeny derived from gamma-irradiation ranging from 0.0 to 2.0 Gy/d were used to identify mutations, including single base substitutions (SBSs) and small insertions/deletions (INDELs). We aimed to evaluate the relationship between genome-scale mutation frequency and the low-to-middle dose of radiation using statistical modeling. Furthermore, we assessed the ratio of heterozygous to homozygous mutations in the M_2_ generation, to gain insight into radiosensitivity in the reproductive stage compared with that in the vegetative stage.

## MATERIALS AND METHODS

### Plant materials, gamma irradiation, and whole-genome sequencing

Detail information of the irradiation experiment is provided in Supplemental Text S1. *Arabidopsis thaliana* L. (Columbia-0) seeds (M1 plants) were germinated in 28-mm diameter pots filled with a perlite-vermiculite mixture (1:1). M_1_ plants were exposed to chronic gamma irradiation throughout the life-cycle—from emergent seedlings (5 days after sowing) until seed maturity at two months—using a ^137^Cs gamma irradiator. Dose rates were set at 0.0, 0.4, 1.4, and 2.0 Gy/d as control, low, middle, and high treatment levels, respectively. Self-fertilized M_2_ seeds were obtained from the irradiated M_1_ plants; three M_2_ plants were derived from each of three M_1_ plants in each of the four treatments, and were grown under control conditions. A total of 48 plants, 12 M_1_ and 36 M_2_ plants, were used for whole-genome sequencing (Supplemental Figure S1). Genomic DNA was extracted using a modified CTAB method and a DNeasy Plant Mini Kit (Qiagen) following the manufacturer’s protocol for the M_1_ and M_2_ plants, respectively. Libraries constructed using TruSeq Nano DNA Kit (Illumina, CA, USA) were sequenced on an Illumina HiSeqX platform, producing 150-bp paired-end reads. The raw reads were deposited in the DDBJ/GenBank/EMBL database under the accession number DRA009784.

### Mutation identification

All custom scripts and commands for computational analyses can be found at https://github.com/akihirao/At_Reseq. Adaptor trimming and quality assessment were performed using fastp (ver. 0.20.0) [24]. Cleaned reads were mapped to the *A. thaliana* TAIR10 reference genome (The Arabidopsis Information Resource at www.arabidopsis.org) using BWA (ver. 0.7.17) [25]. The variant calling procedure was adapted from the germline short variant discovery workflow in GATK (ver. 4.1.7.0) [26]. Variants of SBSs and small INDELs were called across all individuals within the same run of the HaplotypeCaller in GATK. Raw variants were filtered using GATK VariantFiltration with the parameters as described in the custom scripts. Variants in violation of the Mendelian parent–progeny relationship were identified as candidate mutations. The candidate mutations having an allele frequency (AF, proportion of mutant reads at a site) of 80% or more were assigned to homozygous mutations; the candidate mutations with 25% < AF < 80% were assigned to heterozygous mutations; candidates having AF of 25% or less were excluded. Candidate mutations found in more than two M_2_ plants of full-sibs within a family were included as family-shared mutations; those found in more than two plants across families were excluded as background mutations. Candidate INDEL mutations were verified by the split read approach using Pindel (ver. 0.2.5) [27]. Close-positioned mutations within 150 bp of each other in a single sample were excluded, as adjacent mutation candidates include a high proportion of false positives due to mismatch mapping [28]. Therefore, in this study, complex structural mutations of more than two SBSs and/or INDEL mutations within 150 bp were not taken into account. The identified mutation sites were confirmed using IGV (ver. 2.4.13) [29]. The haploid mutation rate was estimated with the equation μ = (*m*/2*n*) × *c*, where μ represents the mutation rate per nucleotide site per generation, *m* is the number of mutations identified in diploid M_2_ plants, *n* is the number of reference nucleotide sites accessible for variant calling, and *c* represents the confirmation rate of identified mutations by the Sanger sequencing platform. A final dataset of mutations was applied for annotation using SnpEff (ver. 4.3) [30] with default settings. The putative deleterious effects of mutations on gene function were classified into the following four categories: high (e.g., frameshift mutations and stop-gain mutations)-, moderate (e.g., missense mutations)-, low (e.g., synonymous mutations)-, and modifier-impact mutations (e.g., intron and intergenic mutations) (see http://snpeff.sourceforge.net for details). Locations of mutations on chromosomes were visualized with Circos (ver. 0.69) [31].

### In silico mutation simulation

To test the accuracy of our mutation calling pipelines, in silico simulation was performed. We simulated 3000 random mutations (2000 SBSs and 1000 INDELs) throughout diploid genome of M_1_ individual using simuG (ver. 1.0.0) [32], and investigated whether our pipeline recovered the simulated mutations in selfed M_2_ progeny. Diploid genome of M_2_ progeny was randomly simulated from that of the M_1_ mutant under the Mendelian rule using the custom script. Thus, all simulated mutations were assumed to occur prior to gametogenesis of M_1_ individual. Simulated genomes of three independent M_1_ mutants and nine M_2_ individuals—three M_2_ individuals derived from each of the three M_1_ mutants—were generated, and then their corresponding raw sequence reads were also simulated using sandy (ver. 0.23) (https://galantelab.github.io/sandy/). Read mapping, variant calling, and mutation identification procedures were followed as mentioned above. All custom scripts and commands for the mutation simulation can be found at https://github.com/akihirao/At_Reseq_sim.

### Verifying mutations by Sanger sequencing platform

Sanger sequencing and fragment size analysis were conducted to validate select SBS and INDEL mutations identified by whole genome sequencing, respectively (see Supplemental Text S2 for detail). The specific primers for 31 randomly chosen SBS mutation sites and 83 INDEL mutation sites—all of 47 INDEL sites on chromosome 1, and 36 randomly chosen INDEL sites on the other chromosomes—were designed by Primer3 (ver. 2.3.7) [33]. Sanger sequences of each of the SBS sites from the M_2_ mutant plants and their maternal/full-sib plants were obtained with ABI3130 and ABI3730 (Applied Biosystem). PCR amplicons of each of the INDEL sites were analyzed for fragment sizes on an ABI3130. Allele-specific fragment sizes were determined using GeneMapper (ver. 4.0) (Applied Biosystems) and GeneMaker (ver. 1.6) software (SoftGenetics, PA, USA).

### Statistical analyses

All statistical analyses were performed using the open source system, R (ver. 4.0.2) [34]. The effects of irradiation on radiobiological end-points (i.e., the number of total mutations and SBS/insertion/deletion mutations; the number of high-, moderate-, low-, and modifier-impact mutations) were assessed using three statistical models: the linear-quadratic (LQ) model, the Poisson regression model, and the negative binominal (NB) model. LQ models have been widely used for analyzing radiobiological end-points [35]. The Poisson and NB models are appropriate for discrete data such as the number of nucleotide mutations and chromosome aberrations [e.g., 36, 37, 38], among which the NB model allows for overdispersion. The LQ model was fitted using non-linear regression model with the stats::nls function in R; the Poisson and NB model were fitted using generalized linear models (GLMs) with the stats::glm and MASS::glm.nb functions in R, respectively. Furthermore, a generalized linear mixed-effect model (GLMM) framework was also implemented for the Poisson model and the NB model to incorporate the random effect of family (i.e., M_1_ line) using the lmer::glmer and lme4::glmer.nb functions in R, respectively. Overdispersion of data was tested using the pscl::odTest function in R. Additionally, the relationship between absolute size of mutation and ionizing radiation was analyzed using two-part hurdle model with Poisson or negative binomial distribution. The hurdle analysis was in two-step process: (1) determining the probability of whether or not INDEL mutations, but not SBS mutations, will occur (2) and, if INDELs, determining the significant drivers of absolute size of mutation. The hurdle analysis was conducted using the pscl::hurdle function in R. Overall, the best-fitting model among the candidate models was selected based on Akaike Information Criterion (AIC). For the ratio of heterozygous to homozygous mutations within individual plants, exact binomial tests were conducted to assess whether the proportion of heterozygous mutations among total mutations was significantly greater than the expected Mendelian segregation ratio.

## RESULTS

### Detection of mutation

Whole genome resequencing of 48 samples—12 M_1_ and 36 M_2_ samples—produced a total of 3493.3 million pair-end reads (527.5 Gbp), with an average of 72.8 million reads (11.0 Gbp) per sample (Supplementary Table S1). After removing the low quality, unpaired, and duplicate reads, more than 99.5% of clean reads were mapped to the TAIR10 reference genome. The average depth of coverage was 65X and 98.4% of the genome on average had at least 10X coverage.

A total of 586 *de novo* mutation sites—400 SBSs and 186 INDELs (171 deletions and 15 insertions)—were identified in the 36 plants in the M_2_ generation (Figure 1 and Supplementary Table S2). Of the 586 mutation sites, 30 mutation sites (23 SBSs and 7 deletions) were identified in multiple plants of full-sibs as family-shared mutations. Our pipeline recovered more than 97% of simulated mutations (Supplementary Table S3). The confirmation rates of identified mutations (*c*) estimated by the Sanger sequencing platform were 93.3% (28/30) and 87.8% (65/74) of the SBSs and INDELs, respectively, while one and nine of the surveyed 31 SBSs and 83 INDELs, respectively, were undetermined because the primers failed to amplify the region (Supplementary Table S4). Thereafter, all of the 586 mutation sites were used for downstream analyses.

**Figure 1.**
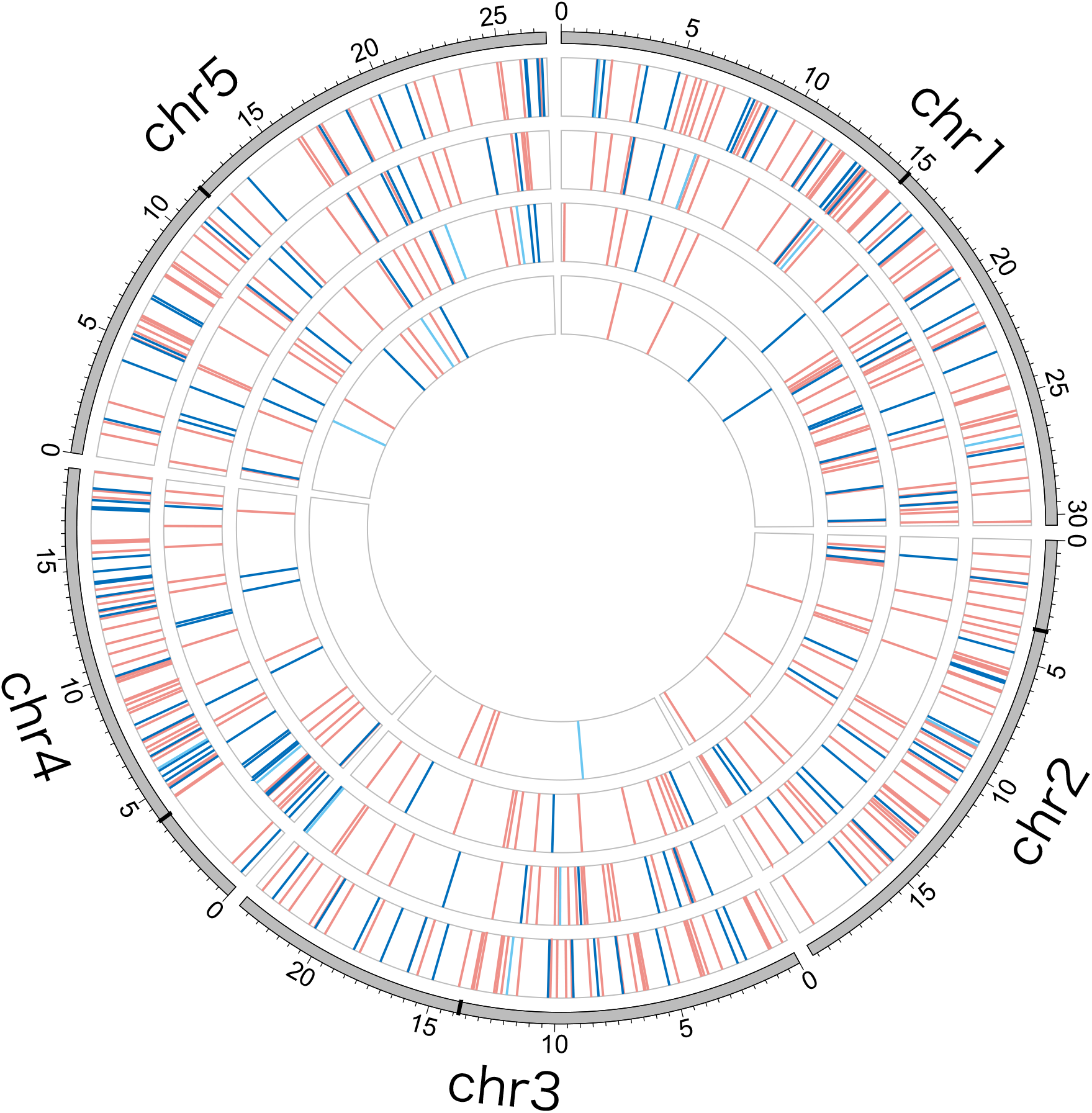
Circos diagram illustrating *de novo* mutations in 36 M_2_ *Arabidopsis thaliana* plants. Mutations in each of the control, low, middle, and high dose irradiated treatments are indicated by short lines on each of the four interior circles from the inner to the outer circles. Pink, blue, and light-blue lines represent single base substitution, deletion, and insertion mutations, respectively.

### The effect of gamma irradiation on mutations

Mutation frequency and mutation rate increased approximately 5- to 14-fold with an increase in the irradiation dose from 0.0 to 2.0 Gy/d (Table 1). The best-fitting statistical model for the relationship between the number of total mutations and ionizing radiation was the NB model in the GLMM (Supplementary Table S5), accounting for the overdispersion of mutation frequency (*p* < 0.001, by the likelihood ratio test). SBSs and deletions, but not insertions, were also significantly increased by irradiation (Figure 2 and Supplementary Table S5). The proportions of SBSs among total mutations were not significantly different among the treatments (*p* > 0.05, by Fisher exact test with Bonferroni correction). The frequency distribution of INDEL size showed that the most frequent type of INDEL was a single base deletion (Supplementary Figure S2). The absolute size of mutation also significantly increased with irradiation (Supplementary Table S6). The transition/transversion (Ti/Tv) ratio was 0.56 ± 0.48, 0.67 ± 0.44, and 0.69 ± 0.23 in the low, middle, and high treatments, respectively. The Ti/Tv ratios for irradiated progeny were significantly different from the Ti/Tv ratio in spontaneous mutations (approximately 0.2- to 0.3-fold lower, estimated from [21]) (*p* < 0.05, by chi-squared test), although the Ti/Tv ratio in the control across samples (6/9 = 0.677) was an inaccurate estimate due to zero-inflated observations, i.e., two out of nine control samples had no SBS mutations.

**Table 1.** Summary of mutation frequency and mutation rate in M_2_ plants of *Arabidopsis thaliana* (mean ± SD).

| Irradiation treatment | No. total mutations | No. SBS mutations | No. deletion mutations | No. insertion mutations | No. family-shared mutations | Mutation rate (/site/generation) |
| --- | --- | --- | --- | --- | --- | --- |
| Control (0.0 Gy/d) | 2.4 ± 1.5 | 1.7 ± 1.4 | 0.4 ± 0.7 | 0.3 ± 0.5 | 0.4 ± 0.9 | 0.95E-08 ± 0.59E-08 |
| Low (0.4 Gy/d) | 12.3 ± 5.3 | 8.4 ± 2.7 | 3.7 ± 2.5 | 0.2 ± 0.4 | 1.3 ± 1.3 | 4.79E-08 ± 2.03E-08 |
| Middle (1.4 Gy/d) | 18.8 ± 3.3 | 12.2 ± 4.0 | 6.0 ± 2.0 | 0.6 ± 0.5 | 1.1 ± 2.0 | 7.29E-08 ± 1.33E-08 |
| High (2.0 Gy/d) | 35.1 ± 10.0 | 24.8 ± 7.0 | 9.8 ± 5.0 | 0.6 ± 0.7 | 4.0 ± 4.7 | 13.68E-08 ± 3.86E-08 |

**Figure 2.**
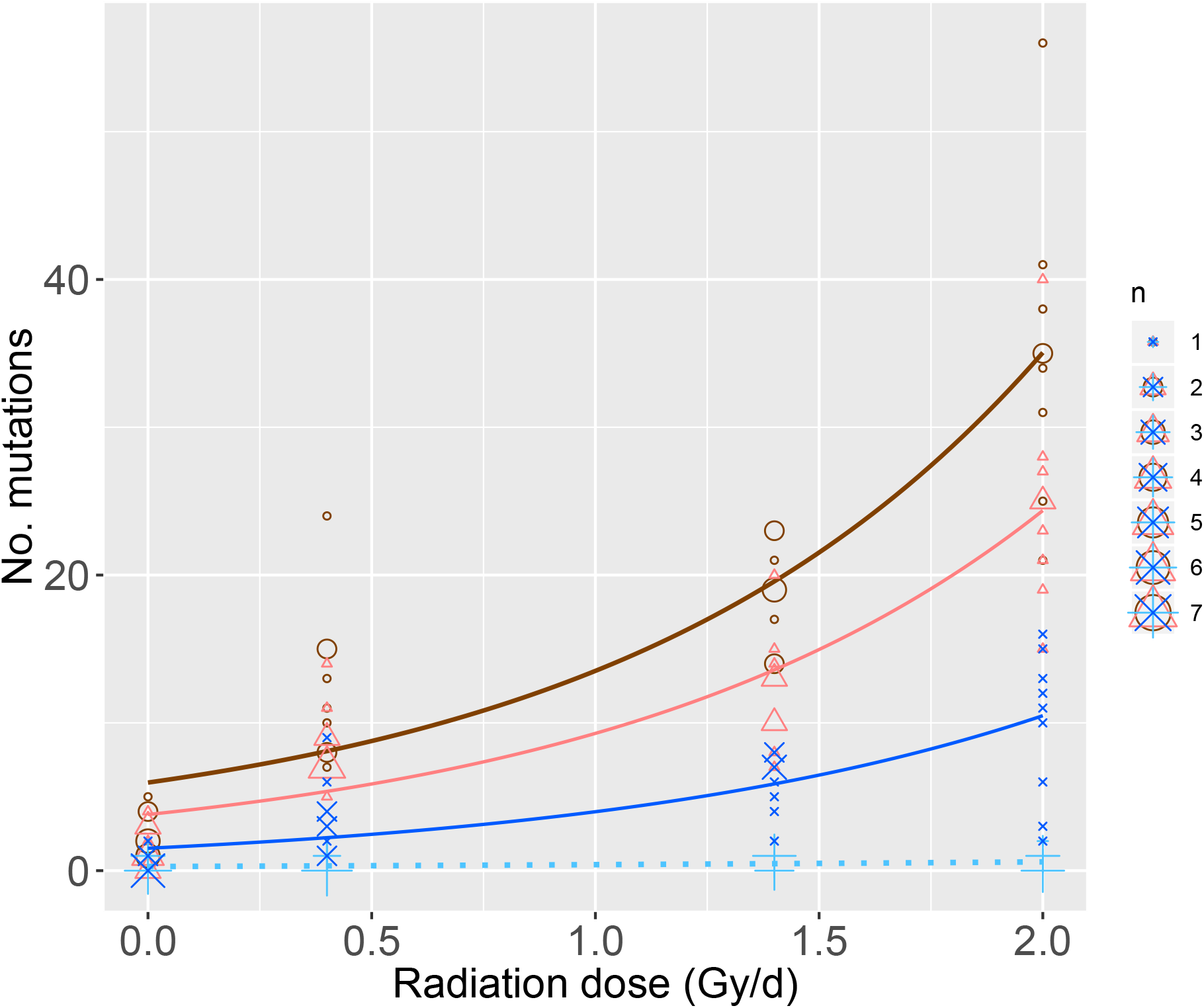
Observed mutation frequency and predicted estimate for the best-fitting negative binominal model in response to gamma irradiation. Open circle, total mutations; open triangles, single base substitutions; crosses, deletions; pluses, insertions. See Supplementary Table S5 for the detailed results of the best-fitting models, which were selected using Akaike’s information criterion.

### Zygosity of mutations

In this study, homozygous mutations found in M_2_ plants must be descended from mutation events that occurred before gamete formation of the M_1_ plants, while heterozygous mutations could be derived from mutation events not only before but also after gametogenesis. The frequencies of heterozygous mutations were excessively higher than those of homozygous mutations in the radiation treatments, especially in the middle and high treatments (Figure 3). Exact binomial tests showed that the proportion of heterozygous mutations among total mutations was significantly greater than the expected Mendelian ratio for all nine M_2_ plants in the high treatment (*p* < 0.05), for eight out of nine plants in the middle treatment (*p* < 0.05), and for four out of nine in the low treatment (*p* < 0.05), but not for any of the nine in the control treatment (*p* > 0.05) (Supplementary Table S7). On the other hand, the in silico mutation simulation assuming all mutations occur prior to gametogenesis of M_1_ generation showed that the ratio of heterozygous to homozygous mutations in M_2_ plants followed the Mendelian segregation ratio (*p* > 0.05, Supplementary Table S3). These results support the hypothesis that mutational effects of ionizing radiation are more intensive after than before gametogenesis for higher dosage irradiated plants.

**Figure 3.**
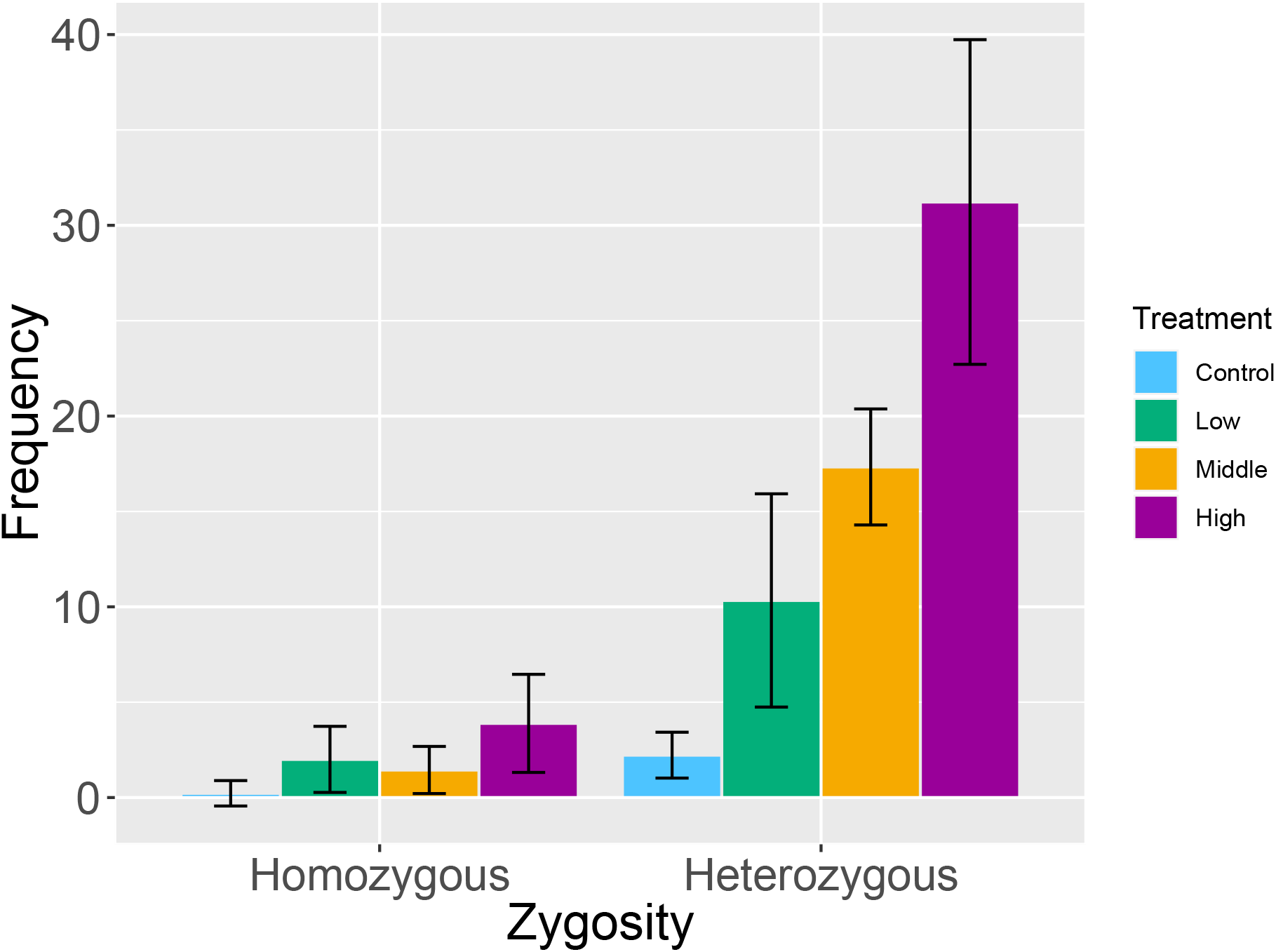
Frequency of homozygous and heterozygous mutations in selfed M_2_ progeny derived from the control (0.0 Gy/d), low (0.4 Gy/d), middle (1.4 Gy/d), and high (2.0 Gy/d) dose irradiated plants (means ± SD).

### Mutation effects on gene function

Of the 586 mutation candidates, 45 (7.7%), 101 (17.2%), 27 (4.6%) and 413 (70.5%) mutations had possible high-, moderate-, low-, and modifier-impacts on gene function, respectively (Supplementary Table S2). The best-fitting statistical model for the relationship between the number of high-impact mutations and irradiation dose showed a significant increasing effect of irradiation (Supplementary Figure S3 and Table S8). The same treads in radiation response were also observed for the number of moderate-, low-, and modifier-impact mutations, respectively (Supplementary Figure S3 and table S8).

## DISCUSSION

Genome-wide mutation frequency increased in response to the low-to-middle dose rates of ubiquitous radiation (lower than 2.0 Gy/d). The increasing pattern of *de novo* mutations in response to the dose rates of irradiation fitted the NB model, indicating that radiation-induced mutation frequency highly varies among samples. Among the types of mutations, SBSs were more prevalent than INDELs, with deletions being more frequent than insertions.

The estimated mutation rate of the control, i.e., spontaneous mutation rate, was 9.5 × 10^−9^ mutations per site per plant generation, which are comparable with the standard and more fine-grained estimates from mutation accumulation lines of *A. thaliana* (7.1 × 10^−9^ and 8.3 × 10^−9^ from [21] and [22], respectively). SBS mutations were the major type of spontaneous mutations (71% of the total), in agreement with previous research (7.0 × 10^−9^ and 1.3 × 10^−9^ for SBS and INDEL, respectively [22]). Mutation frequency increased several- to a dozen-fold in response to the dose rate of irradiation, while the major proportion of SBS was constant; however, the mutation spectrum was changed. It is worth noting that the mutation frequency was observed to vary greatly in each plant, even if the random effect of family M_1_ line was substantial, as shown by that the best-fitting models accounting for overdispersion. In general, transitions are major spontaneous mutations rather than transversions (the Ti/Tv ratio: ~3) [21, 22]. The increased frequencies of transversion for the irradiated progeny (the Ti/Tv ratios = 0.6– 0.7) were compatible with previous estimates from plants that were exposed to various types of ionizing irradiation [6, 9, 10, 12–14]. Additionally, absolute size of mutations increased in response to dose of irradiation.

Of particular note is that INDEL mutations found in this study were limited to relatively small sizes, because Illumina short-read sequencing technology combined with current tools for INDEL detection are still far from identifying the complete list of INDEL mutations, especially those of a larger nature (see [8] for more detailed discussion). The observed sizes of INDEL mutations were less than 116 bp (Supplementary Table S2 and Figure S2). Naito et al. (2005) [39] showed that irradiation with gamma-rays (150-600 Gy) or carbon ions (40-150 Gy) to dry pollen in *Arabidopsis* mainly induced large fragment deletions of up to > 6 Mbp in the M_2_ progeny, although the majority of the large INDELs could not be transmitted to M_3_ progeny. Life-cycle chronic irradiation could act directly on pollen of the M_1_ plants. Thus, the possibility of extremely large structural variants and the mutation load remain as further directions, for which long-read platforms and/or the long-insert paired-end libraries may be much more suitable.

The observed number of heterozygous mutations were significantly greater that the expected Mendelian segregation ratios for more highly irradiated samples, suggesting that the mutational effects of life-cycle chronic radiation are more intensive in the reproductive stage. In this study system, homozygous mutations found in selfed M_2_ progeny must be descended from *de novo* mutation events before gamete formation in M_1_ plants, while heterozygous mutations could be derived from mutation events not only before but also after gametogenesis. It is unlikely that the highly excessive number of heterozygous mutations resulted from purging homozygosity of mutations with recessive lethal effects, because the annotation analysis shows that high-impact mutations (i.e., loss-of-function gene mutations) accounted for only 7.7% of total mutations (Supplementary Table S2). Previous studies of the genetic consequences of acute/chronic radiation only during the vegetative stage showed that the heterozygous/homozygous ratio of mutations followed the Mendelian segregation ratio [e.g., 6, 12, 14, 40]. On the other hand, in this study, life-cycle chronic irradiation induced a greater number of heterogeneous mutations that should occur after gametogenesis, i.e., any reproductive stage from gamete formation to fertilization, zygote development, and then seed maturation. Therefore, we concluded that mutational radiosensitivity is higher in the reproductive stage, especially for more irradiated plants.

From the other point of view, family-shared mutations (i.e., multiple common mutations within a full-sib family) were interpreted as evidence for mutation events that occurred prior to gametogenesis. According to the lows of Mendelian inheritance, if *de novo* germline mutations were occurred in the vegetative growth stage, 84.4% of these mutations would be transmitted to more than two out of the three full-sib M_2_ progeny and then assigned as family-shared mutations in this study system. However, the observed percentages of family-shared mutations in total mutations were low for the irradiated progeny (6.3-11.8%: Table 1 and Supplementary Table S2). In a strict sense, family-shared mutations were derived from mutation events not only in the vegetative growth stage but also in the early reproductive stage, after bolting until gametogenesis. Thus, low percentages of family-shared mutations for the irradiated progeny, even if possibly including some mutation events in the early reproductive stage, support at least the hypothesis that radiation-induced mutations occurred less frequently in the vegetative stage.

Mutations occur unevenly across the developmental period. The idea of elevated mutagenicity in meiosis stems from an early 1960s study of Magni and Von Bostel [41], in which yeast *Saccharomyces cerevisiae* demonstrated a higher mutation rate in meiosis than in mitosis. This phenomenon, of elevated levels of mutation in meiosis, was then observed in other organisms such as mice, and further extended from several loci to genome-scale assessments [reviewed in 42]. Thus, our results may suggest elevated radiosensitivity especially in meiosis, or within a chronologically close period, as mentioned below. A pollen irradiation experiment in *Arabidopsis* showed that meiosis and earlier developmental stages of pollen were the most irradiation-sensitive stage for fertility, while high frequencies of targeted mutations were obtained by irradiation from the second mitotic division of pollen grains to the mature pollen stages [43]. Moreover, haploid phases such as pollen and embryo sacs are expected to reduce the opportunity for conservative DNA repair involving homologous chromosomes, thus increasing the mutation rate. Furthermore, pollen, as male gamete, might be more radiosensitive than the female gamete. Further investigation with cross-pollination experiments between irradiated and non-irradiated plants will be necessary to clarify radiosensitivity in male and female gametes.

In conclusion, this study revealed the mutation profile and frequency of SBS and small INDEL mutations induced by chronic irradiation in *A. thaliana* at the whole genome level. Increasing mutations in response to the dose rate of irradiation showed that mutation frequency is highly variable in its character. Furthermore, we observed that the mutational effects of life-cycle chronic radiation are more intensive in the reproductive stage. These outcomes could provide valuable clues for practical strategies for the environmental radioprotection of plants both on Earth and in space.

## Data accessibility

Raw FASTQ files were deposited in the DRA/SRA/ERA under the accession number DRA009784. All the code necessary to reproduce this analysis can be accessed from: https://github.com/akihirao/At_Reseq and https://github.com/akihirao/At_Reseq_sim.

## Authors contributions

S.K., Y.W., S.U. and A.S.H. designed the study; Y. W. was performed the irradiation experiment; A.S.H., Y.H. and T.T. were performed the molecular laboratory experiments; A.S.H. conducted the data analyses and wrote the manuscript with contributions from the co-authors.

## Acknowledgments

We are grateful to Y. Hase and K. Satoh for the information on the analysis pipeline they so kindly supplied. We also thank H. Yasuda, H. Setoguchi, Y. Isagi, T, and Y. Tsumura for valuable comments on this work.

## REFERENCES

[1] Muller, H.J. 1927 Artificial transmutation of the gene. Science 66, 84–87.

[2] Stadler, L.J. 1928 Genetic effects of X-rays in maize. Proc. Natl. Acad. Sci. USA 14, 69.

[3] Mousseau, T.A. & Moller, A.P. 2020 Plants in the light of ionizing radiation: What have we learned From Chernobyl, Fukushima, and other “hot” places? Front Plant Sci 11, 552. (doi:10.3389/fpls.2020.00552).

[4] Caplin, N. & Willey, N. 2018 Ionizing radiation, higher plants, and radioprotection: from acute high doses to chronic low doses. Front Plant Sci 9, 847. (doi:10.3389/fpls.2018.00847).

[5] Nikitaki, Z., Hola, M., Dona, M., Pavlopoulou, A., Michalopoulos, I., Angelis, K.J., Georgakilas, A.G., Macovei, A. & Balestrazzi, A. 2018 Integrating plant and animal biology for the search of novel DNA damage biomarkers. Mutat. Res. 775, 21–38. (doi:10.1016/j.mrrev.2018.01.001).

[6] Belfield, E.J., Gan, X., Mithani, A., Brown, C., Jiang, C., Franklin, K., Alvey, E., Wibowo, A., Jung, M., Bailey, K., et al. 2012 Genome-wide analysis of mutations in mutant lineages selected following fast-neutron irradiation mutagenesis of *Arabidopsis thaliana*. Genome Res. 22, 1306–1315. (doi:10.1101/gr.131474.111).

[7] Hirano, T., Kazama, Y., Ishii, K., Ohbu, S., Shirakawa, Y. & Abe, T. 2015 Comprehensive identification of mutations induced by heavy-ion beam irradiation in Arabidopsis thaliana. Plant J. 82, 93–104. (doi:10.1111/tpj.12793).

[8] Shirasawa, K., Hirakawa, H., Nunome, T., Tabata, S. & Isobe, S. 2016 Genome-wide survey of artificial mutations induced by ethyl methanesulfonate and gamma rays in tomato. Plant Biotechnol. J. 14, 51–60. (doi:10.1111/pbi.12348).

[9] Du, Y., Luo, S., Li, X., Yang, J., Cui, T., Li, W., Yu, L., Feng, H., Chen, Y., Mu, J., et al. 2017 Identification of substitutions and small insertion-deletions induced by carbon-ion beam irradiation in *Arabidopsis thaliana*. Front Plant Sci 8, 1851. (doi:10.3389/fpls.2017.01851).

[10] Kazama, Y., Ishii, K., Hirano, T., Wakana, T., Yamada, M., Ohbu, S. & Abe, T. 2017 Different mutational function of low- and high-linear energy transfer heavy-ion irradiation demonstrated by whole-genome resequencing of *Arabidopsis* mutants. Plant J. 92, 1020–1030. (doi:10.1111/tpj.13738).

[11] Du, Y., Luo, S., Yu, L., Cui, T., Chen, X., Yang, J., Li, X., Li, W., Wang, J. & Zhou, L. 2018 Strategies for identification of mutations induced by carbon-ion beam irradiation in Arabidopsis thaliana by whole genome re-sequencing. Mutat. Res. 807, 21–30. (doi:10.1016/j.mrfmmm.2017.12.001).

[12] Hase, Y., Satoh, K., Kitamura, S. & Oono, Y. 2018 Physiological status of plant tissue affects the frequency and types of mutations induced by carbon-ion irradiation in *Arabidopsis*. Sci Rep 8, 1394. (doi:10.1038/s41598-018-19278-1).

[13] Li, F., Shimizu, A., Nishio, T., Tsutsumi, N. & Kato, H. 2019 Comparison and characterization of mutations induced by gamma-ray and carbon-ion irradiation in rice (*Oryza sativa* L.) using whole-genome resequencing. G3 9, 3743–3751. (doi:10.1534/g3.119.400555).

[14] Hase, Y., Satoh, K., Seito, H. & Oono, Y. 2020 Genetic consequences of acute/chronic gamma and carbon ion irradiation of *Arabidopsis thaliana*. Front Plant Sci 11. (doi:10.3389/fpls.2020.00336).

[15] Jo, Y.D. & Kim, J.-B. 2019 Frequency and spectrum of radiation-induced mutations revealed by whole-genome sequencing analyses of plants. Quantum Beam Science 3, 7.

[16] United Nations Scientific Committee on the Effects of Atomic Radiation (UNSCEAR). 2000 Sources and effects of ionizing radiation. In: Report to the General Assembly, with Scientific Annexes. United Nations, New York.

[17] Kodaira, S., Tolochek, R.V., Ambrozova, I., Kawashima, H., Yasuda, N., Kurano, M., Kitamura, H., Uchihori, Y., Kobayashi, I., Hakamada, H., et al. 2014 Verification of shielding effect by the water-filled materials for space radiation in the International Space Station using passive dosimeters. Advances in Space Research 53, 1–7. (doi:10.1016/j.asr.2013.10.018).

[18] McLean, A.R., Adlen, E.K., Cardis, E., Elliott, A., Goodhead, D.T., Harms-Ringdahl, M., Hendry, J.H., Hoskin, P., Jeggo, P.A., Mackay, D.J.C., et al. 2017 A restatement of the natural science evidence base concerning the health effects of low-level ionizing radiation. Proc R Soc B 284, 20171070. (doi:10.1098/rspb.2017.1070).

[19] United Nations Scientific Committee on the Effects of Atomic Radiation (UNSCEAR). 1996 Sources and effects of ionizing radiation. In: Report to the General Assembly, with Scientific Annexes. United Nations, New York.

[20] International Commission of Radiological Protection (ICRP). 2014 Protection of the environment under different exposure situations. In: ICRP Publication 124. Ann. ICRP 43.

[21] Ossowski, S., Schneeberger, K., Lucas-Lledo, J.I., Warthmann, N., Clark, R.M., Shaw, R.G., Weigel, D. & Lynch, M. 2010 The rate and molecular spectrum of spontaneous mutations in *Arabidopsis thaliana*. Science 327, 92–94. (doi:10.1126/science.1180677).

[22] Weng, M.L., Becker, C., Hildebrandt, J., Neumann, M., Rutter, M.T., Shaw, R.G., Weigel, D. & Fenster, C.B. 2019 Fine-grained analysis of spontaneous mutation spectrum and frequency in *Arabidopsis thaliana*. Genetics 211, 703–714. (doi:10.1534/genetics.118.301721).

[23] Halligan, D.L. & Keightley, P.D. 2009 Spontaneous mutation accumulation studies in evolutionary genetics. Annu. Rev. Ecol. Evol. Syst. 40, 151–172. (doi:10.1146/annurev.ecolsys.39.110707.173437).

[24] Chen, S., Zhou, Y., Chen, Y. & Gu, J. 2018 fastp: an ultra-fast all-in-one FASTQ preprocessor. Bioinformatics 34, i884–i890. (doi:10.1093/bioinformatics/bty560).

[25] Li, H. & Durbin, R. 2009 Fast and accurate short read alignment with Burrows-Wheeler transform. Bioinformatics 25, 1754–1760. (doi:10.1093/bioinformatics/btp324).

[26] McKenna, A., Hanna, M., Banks, E., Sivachenko, A., Cibulskis, K., Kernytsky, A., Garimella, K., Altshuler, D., Gabriel, S. & Daly, M. 2010 The Genome Analysis Toolkit: a MapReduce framework for analyzing next-generation DNA sequencing data. Genome Res. 20, 1297–1303. (doi:10.1101/gr.107524.110).

[27] Ye, K., Schulz, M.H., Long, Q., Apweiler, R. & Ning, Z. 2009 Pindel: a pattern growth approach to detect break points of large deletions and medium sized insertions from paired-end short reads. Bioinformatics 25, 2865–2871. (doi:10.1093/bioinformatics/btp394).

[28] Keightley, P.D., Ness, R.W., Halligan, D.L. & Haddrill, P.R. 2014 Estimation of the spontaneous mutation rate per nucleotide site in a *Drosophila melanogaster* full-sib family. Genetics 196, 313–320. (doi:10.1534/genetics.113.158758).

[29] Thorvaldsdóttir, H., Robinson, J.T. & Mesirov, J.P. 2013 Integrative Genomics Viewer (IGV): high-performance genomics data visualization and exploration. Briefings in bioinformatics 14, 178–192.

[30] Cingolani, P., Platts, A., Wang, L.L., Coon, M., Nguyen, T., Wang, L., Land, S.J., Lu, X. & Ruden, D.M. 2012 A program for annotating and predicting the effects of single nucleotide polymorphisms, SnpEff: SNPs in the genome of *Drosophila melanogaster* strain w1118; iso-2; iso-3. Fly 6, 80–92. (doi:10.4161/fly.19695).

[31] Krzywinski, M., Schein, J., Birol, I., Connors, J., Gascoyne, R., Horsman, D., Jones, S.J. & Marra, M.A. 2009 Circos: an information aesthetic for comparative genomics. Genome Res. 19, 1639–1645. (doi:10.1101/gr.092759.109).

[32] Yue, J.X. & Liti, G. 2019 simuG: a general-purpose genome simulator. Bioinformatics 35, 4442–4444. (doi:10.1093/bioinformatics/btz424).

[33] Untergasser, A., Cutcutache, I., Koressaar, T., Ye, J., Faircloth, B.C., Remm, M. & Rozen, S.G. 2012 Primer3—new capabilities and interfaces. Nucleic Acids Res. 40, e115–e115. (doi:10.1093/nar/gks596).

[34] R Core Team. 2020 R: A language and environment for statistical computing. In R Core Team (Vienna, Austria, R Foundation for Statistical Computing.

[35] Sachs, R.K. & Brenner, D.J. 1998 The mechanistic basis of the linear-quadratic formalism. Medical physics 25, 2071–2073.

[36] Brame, R.S. & Groer, P.G. 2002 Bayesian analysis of overdispersed chromosome aberration data with the negative binomial model. Radiat. Prot. Dosimet. 102, 115–119.

[37] Edwards, A., Lloyd, D. & Purrott, R. 1979 Radiation induced chromosome aberrations and the Poisson distribution. Radiat. Environ. Biophys. 16, 89–100.

[38] Shuryak, I., Loucas, B.D. & Cornforth, M.N. 2017 Straightening beta: Overdispersion of lethal chromosome aberrations following radiotherapeutic doses leads to terminal linearity in the alpha–beta model. Frontiers in Oncology 7, 318. (doi:10.3389/fonc.2017.00318).

[39] Naito, K., Kusaba, M., Shikazono, N., Takano, T., Tanaka, A., Tanisaka, T. & Nishimura, M. 2005 Transmissible and nontransmissible mutations induced by irradiating *Arabidopsis thaliana* pollen with gamma-rays and carbon ions. Genetics 169, 881–889. (doi:10.1534/genetics.104.033654).

[40] Du, Y., Hase, Y., Satoh, K. & Shikazono, N. 2020 Characterization of gamma irradiation-induced mutations in Arabidopsis mutants deficient in non-homologous end joining. J. Radiat. Res. 61, 639–647. (doi:10.1093/jrr/rraa059).

[41] Magni, G.E. & Von Borstel, R.C. 1962 Different rates of spontaneous mutation during mitosis and meiosis in yeast. Genetics 47, 1097–1108.

[42] Arbel-Eden, A. & Simchen, G. 2019 Elevated mutagenicity in meiosis and its mechanism. Bioessays 41, e1800235. (doi:10.1002/bies.201800235).

[43] Yang, C., Mulligan, B.J. & Wilson, Z.A. 2004 Molecular genetic analysis of pollen irradiation mutagenesis in *Arabidopsis*. New Phytol. 164, 279–288. (doi:10.1111/j.1469-8137.2004.01182.x).

