## Supplemental Figure S1 for "Mutational effects of ubiquitously present gamma radiation on *Arabidopsis thaliana*: insight into radiosensitivity in the reproductive stage"

**Supplementary Figure S1.** Experimental design and plant materials for whole-genome resequencing (WGR). A total of 48 plants, 12 $M_1$ and 36 $M_2$ plants, were used for WGR.

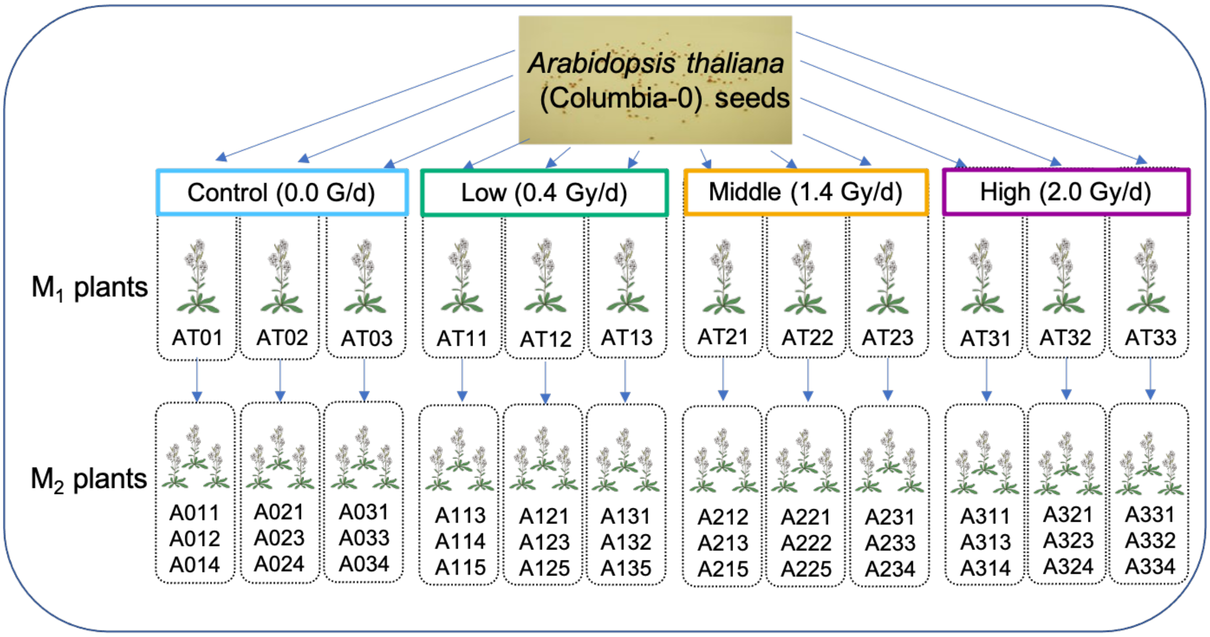


The image of *Arabidopsis* plant is from TogoTV (© 2016 DBCLS TogoTV)
