## Supplemental Figure S2 for "Mutational effects of ubiquitously present gamma radiation on *Arabidopsis thaliana*: insight into radiosensitivity in the reproductive stage"

**Supplementary Figure S2.** Distribution of the size of mutations in the control (0.0 Gy/d), low (0.4 Gy/d), middle (1.4 Gy/d), and high (2.0 Gy/d) dose irradiation treatments, respectively. Insertion and deletion mutations were represented as positive and negative peaks, respectively. Single-base substitution (SBS) mutations are not represented in this figure.

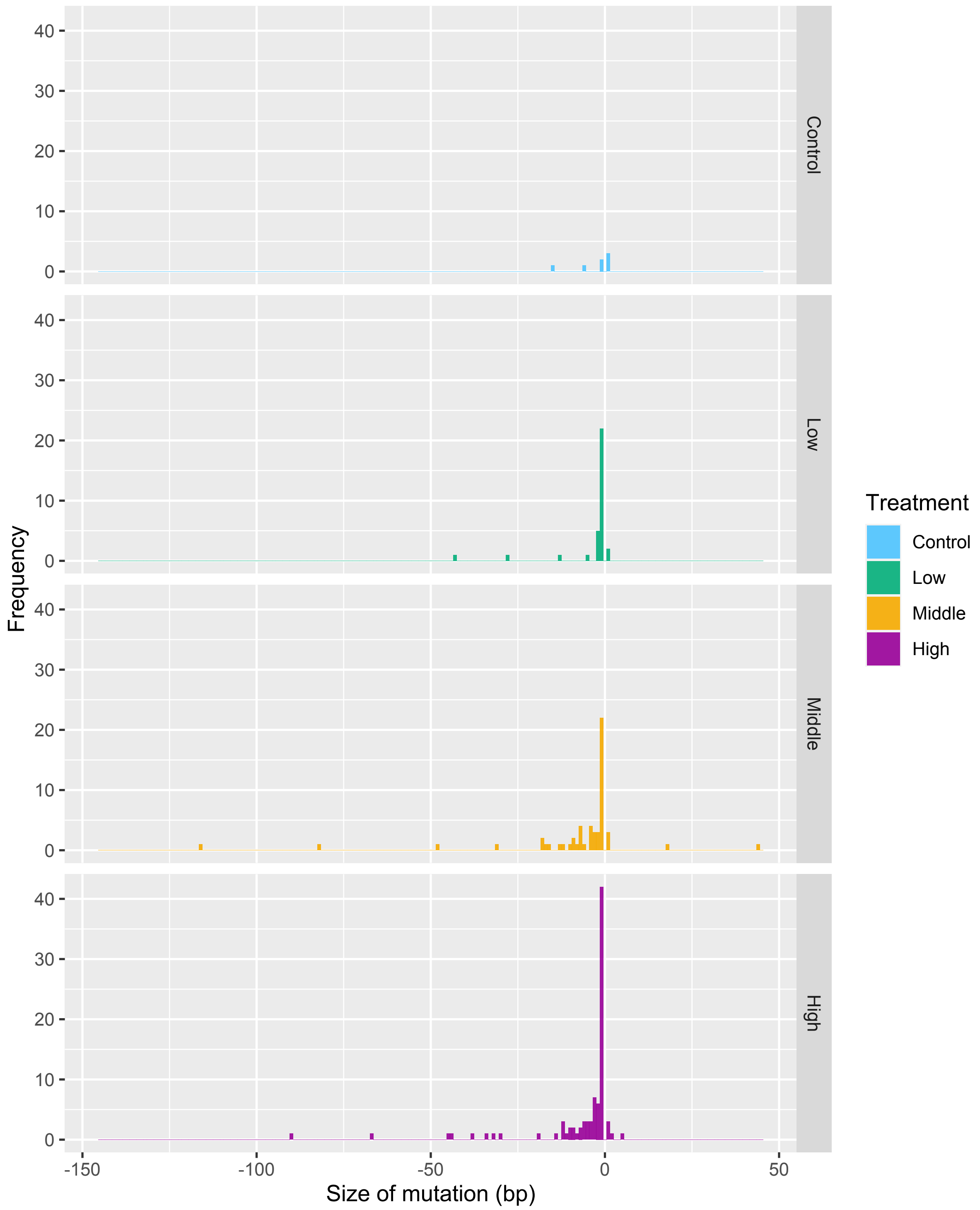
