## Supplemental Figure S3 for "Mutational effects of ubiquitously present gamma radiation on *Arabidopsis thaliana*: insight into radiosensitivity in the reproductive stage"

**Supplementary Figure S3.** Observed mutation frequency in each functional annotation, and predicted estimate for the best-fitting statistical model in response to gamma irradiation: a) high-impact mutations (e.g., frameshift mutations and stop-gain mutations); b) moderate-impact mutations (e.g., missense mutations); c) low-impact mutations (e.g., synonymous mutations); and d) modifier-impact mutations (e.g., intron and intergenic mutations). See Supplementary Table S8 for the detailed results of the best-fitting models, which were selected using Akaike's information criterion.

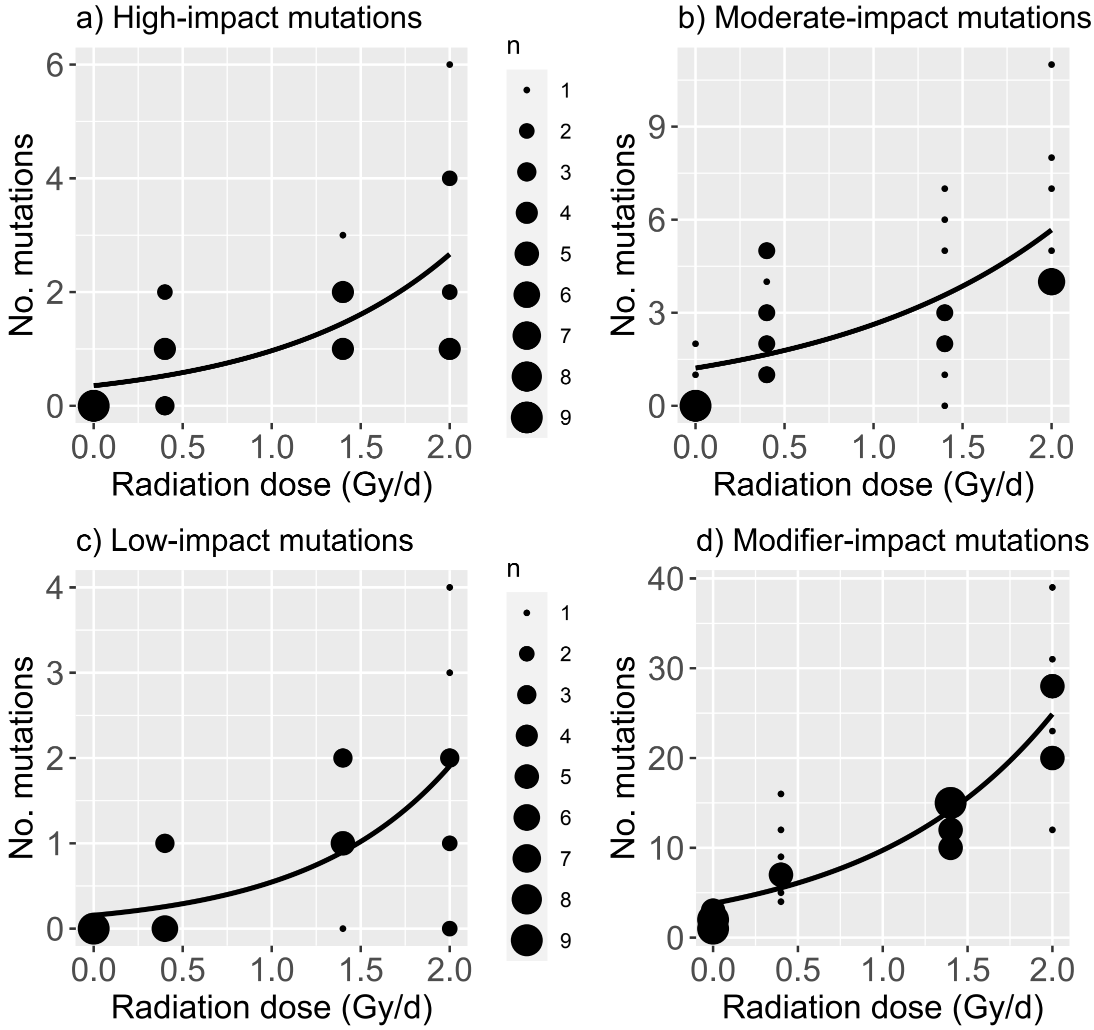
