## Supplemental Text S1 for "Mutational effects of ubiquitously present gamma radiation on *Arabidopsis thaliana*: insight into radiosensitivity in the reproductive stage"

**Supplemental Text S1. Plant materials and gamma irradiation**

*Arabidopsis thaliana* L. (Columbia-0) seeds were obtained from Funakoshi Co., Ltd. (Tokyo, Japan). The dry seeds ($M_1$ plants) were germinated in small pots (28 mm φ × 53 mm H) filled with a 1:1 mixture of perlite and vermiculite. $M_1$ plants were exposed to chronic gamma irradiation throughout the life-cycle—from emergent seedlings (5 days after sowing) until seed maturity at two months—using $^{137}Cs$ gamma irradiator. The $M_1$ plants were placed at specific distances from 7.4 TBq of a $^{137}Cs$ source, where absorbed dose rates were measured using glass rod dosimeters (GD-352M, Chiyoda Technol Corporation, Tokyo, Japan). Dose rates were set at 0.0, 0.4, 1.4, and 2.0 Gy/d as control, low, middle, and high treatment levels, respectively. Self-fertilized $M_2$ seeds were obtained from the irradiated $M_1$ plants. The $M_2$ seeds were surface-sterilized with 0.5% sodium hypochlorite, sown on 1 % agar medium supplemented with 0.5 strength of Murashige and Skoog salt and 2 % of sucrose, and grown for 3 weeks. Growth condition of the $M_1$ and $M_2$ plants were at 22°C under cool-white lamps at an intensity of 100–150 μmol $m^{-2}$ $s^{-1}$ at 16 h/8 h day/night regime. Three $M_2$ plants were derived from each of three $M_1$ plants in each of the four treatments, and were grown under control conditions. A total of 48 plants, 12 $M_1$ and 36 $M_2$ plants, were used for whole-genome sequencing.
