## Supplemental Text S2 for "Mutational effects of ubiquitously present gamma radiation on *Arabidopsis thaliana*: insight into radiosensitivity in the reproductive stage"

**Supplemental Text S2. Checking identified mutations by Sanger sequencer**

Sanger sequencing and fragment size analysis were conducted to validate a part of single base substitution (SBS) and insertion/deletion (INDEL) mutations identified by whole genome sequencing, respectively. The specific primers for 32 SBS mutation sites randomly chosen and 78 INDEL mutation sites—almost all of 48 sites on chromosome 1 and 30 sites randomly chosen from the other chromosomes—were designed by Primer3.

For the Sanger sequencing of the SBS mutation sites, PCRs were carried out in a 10μL total volume with 1μL of template DNA, 2.5 units KOD FX DNA Polymerase (TOYOBO), 1× PCR Buffer for KOD FX (TOYOBO), 0.4 mM each dNTP, and each pair of primers at 0.3 μM. The amplification parameters were 94°C for 2 min; 30 cycles at 98°C for 10 sec, 59°C for 30 sec, and 68°C for 1 min; and finally an elongation step at 68°C for 7 min. The PCR products were purified using an ExoProStar PCR clean-up kit 0.1× (GE Healthcare). The cycle sequencing reactions of forward and reverse strands were performed with the same PCR primers, using 0.1× diluted BigDye v.3.1 (Applied Biosystems). The sequencing products were analyzed on an ABI 3130 and ABI3730 (Applied Biosystems).

For fragment size analysis of the INDEL mutation sites, PCR amplifications with fluorescent dye-labeled primers was performed using a protocol described by Blacket, Robin, Good, Lee, and Miller (2012). PCR amplicons were generated in 10μL total volume with 1μL of template DNA, 5μL of Multiplex PCR Master Mix (QIAGEN), 0.1 μM forward primer, 0.2 μM reverse primer, and 0.1 μM fluorescent dye-labeled primer. The amplification parameters were 95°C for 15 min; 45 cycles at 95°C for 10 sec, 58°C for 90 sec, and 72°C for 1 min; and finally an elongation step at 60°C for 30 min. Allele-specific fragment sizes of PCR amplicons were determined using the GeneMapper ver. 4.0 (Applied Biosystems) and GeneMaker ver. 1.6 (SoftGenetics, PA, USA).

**References**

Blacket, M. J., Robin, C., Good, R. T., Lee, S. F., & Miller, A. D. (2012). Universal primers for fluorescent labelling of PCR fragments--an efficient and cost-effective approach to genotyping by fluorescence. *Molecular Ecology Resources, 12*(3), 456-463. doi:10.1111/j.1755-0998.2011.03104.x
